# CD8 T-cell dysfunction is linked with CAR T-cell failure and can be mitigated by a non-alpha IL-2 agonist, pegenzileukin

**DOI:** 10.1101/2023.08.28.555127

**Authors:** Patrick K Reville, Irtiza N Sheikh, Ahyun Choi, Enyu Dai, Jared Henderson, Xubin Li, Estela Rojas, Cuong Le, Chizitara Okwuchi, Mikielia Devonish, Roberto Carrio, Nathan Pate, Katie Malley, Dinesh Bangari, Julie-Ann Givigan, Chaomei Shi, Bing Liu, Tony Byers, Jason Westin, Sairah Ahmed, Nathan Fowler, Luis Fayad, Hun Ju Lee, Loretta Nastoupil, Ingrid Sassoon, Margot Cucchetti, Rui Wang, Maria Agarwal, Giovanni Abbadessa, Robin Meng, Elamaran Meibalan, Laura Powers, James Cao, Xiaoyou Ying, Kelly Balko, Qunyan Yu, Jing Jiao, Virna Cortez-Retamozo, Sukhvinder Sidhu, Donald Shaffer, Sattva Neelapu, Linghua Wang, Xiangming Li, Michael Green

**Author notes:** Co-first authors (Patrick K Reville, Irtiza N Sheikh, Ahyun Choi, Enyu Dai). Co-senior and corresponding authors (Linghua Wang, Xiangming Li, Michael Green). Correspondence: Michael R. Green, PhD, 1515 Holcombe Blvd., Unit 903, Houston, TX 77030, Xiangming Li, PhD, 350 Water Street, 10FL, Cambridge, MA 02142.

## Abstract

Chimeric antigen receptor (CAR) T-cell therapy has been a breakthrough for relapsed or refractory large B-cell lymphoma (rrLBCL). However, suboptimal CAR T-cell activity can lead to therapeutic failure and dismal outcome. Using single cell RNA-sequencing of rrLBCL tumors, we identify a prominent population of clonally expanded dysfunctional CAR+ CD8 T-cells indicative of ongoing tumor cell engagement, proliferation, and dysfunction at the time of progression from CAR T-cell therapy. Furthermore, we show that rrLBCL patient-derived CAR T-cells are more prone to dysfunction and loss of cytotoxicity compared to healthy donor-derived CAR T-cells. Using both antigen-driven and CAR-driven models of T-cell dysfunction, we show that pegenzileukin, a non-alpha IL2 agonist, can prevent T-cell dysfunction. In both *in vitro* and *in vivo* CAR T-cell models, pegenzileukin improved T-cell expansion and tumor control. This provides pre-clinical rational for use of pegenzileukin in combatting T-cell dysfunction, a central mechanism of CAR T-cell failure.

**HIGHLIGHTS:**

- Tumor-infiltrating CD8 CAR T-cells show clonal expansion and dysfunction at the time of progression.
- rrLBCL patient-derived CAR T-cells are more prone to dysfunction compared to healthy-donor-derived CAR T-cells.
- Pegenzileukin, a non-alpha IL2 agonist, rescues antigen– and CAR-driven CD8 T-cell dysfunction and improves CAR T-cell responses in vivo.

## INTRODUCTION

Autologous CD19-directed chimeric antigen receptor (CAR) T-cell therapy has significantly improved the outcomes of relapsed or refractory large B-cell lymphoma (rrLBCL) compared to standard of care salvage chemoimmunotherapy^1^. However, most patients do not have durable benefit and post-CAR T-cell relapses are difficult to salvage^2^. While the determinants of response to CAR T-cell therapy are multifactorial, one major component relates to T-cell functionality and associated capacity to expand *in vivo*^3^. A higher peak CAR T-cell expansion is significantly associated with durable response for axicabtagene ciloleucel (axi-cel)^4^ and is linked with characteristics of the CAR T-cell infusion product such as the relative frequencies of memory or exhausted CD8 T-cells^5,6^. The development of dysfunctional/exhausted T-cells is driven by a process of repeated and prolonged antigen stimulation and is characterized by decreased proliferation and loss of effector function^7–9^. T-cell dysfunction is associated with decreased tumor cytotoxicity, highlighting the therapeutic rescue of T-cell dysfunction as a potential path towards sustained T-cell proliferation and effector function^10,11^ and more durable responses to CAR T-cell therapy.

To counter T-cell dysfunction, cytokine therapy including interleukin-2 (IL-2) has become an established method shown to stimulate the growth of T-cells and promote their infiltration within tumor, leading to robust tumor cytotoxicity^12,13^. However, while IL-2 is a potent stimulator of T-cell proliferation and differentiation, it also carries clinically significant on-target effects including regulatory T-cells (T_Reg_) expansion and life-threatening vascular leak syndrome due to its stimulation IL-2Rα found on innate lymphoid cells and vascular endothelial tissue^14–16^. T_Reg_ promote T-cell dysfunction and suppress CD8 T-cell effector function and have been implicated in CAR T-cell failure^17,18^. Pegenzileukin (previously THOR-707) is a PEGylated non-alpha IL-2 compound which utilizes site-specific PEGylation to disrupt binding of IL-2Rα while maintaining near-native binding to the IL-2Rβγ dimeric receptor^19^. This allows for amplification of memory CD8 T-cells without T_Reg_ expansion^19^. However, whether non-alpha IL-2 agonists can rescue CAR T-cells from dysfunctional states has not yet been investigated.

Here, we show using single cell RNA-sequencing (scRNA-seq) of primary rrLBCL tumors that post-CAR T-cell relapses have an over-representation of clonally expanded and dysfunctional CD8 CAR T-cells and higher frequencies of T_Reg_ cells. Using *in vitro* models of antigen-driven and CAR-driven T-cell dysfunction, we show that pegenzileukin can prevent or rescuing T-cell dysfunction and driving robust CD8 T-cell proliferation and effector function. In both *in vitro* and *in vivo* CAR T-cell models, the addition of pegenzileukin to CAR T-cells significantly improved tumor clearance. This supports a model in which intratumoral CAR T-cell dysfunction is directly linked with disease progression and may be targeted using non-alpha IL2 agonists such as pegenzileukin.

## RESULTS

### Post-CAR T-cell relapse in rrLBCL shows enrichment of clonally expanded dysfunctional CD8 CAR T-cells

To uncover characteristics associated with CAR T refractoriness, we performed a discovery analysis comparing the tumor microenvironment of rrLBCL tumors from patients that were CAR T naïve (never previously received cell therapy; n=5) compared to CAR T relapsed (progression biopsy obtained post-CAR T therapy, n=7). The average number of preceding lines of therapy (Mann Whitney P-value = 0.3346) and SUV-max of the target lesions (Mann Whitney P-value > 0.99) were not significantly different between groups (Mann Whitney P-value = 0.3346), and the average time to biopsy post CAR T in the refractory group was 120 days (range 48-224) (Table S1). All core needle biopsies were disaggregated into single cell suspensions and run fresh (without prior freezing) on a 10X Chromium with 5’GEX plus TCR and BCR sequencing as previously described^20^. To increase the cell number and associated resolution of cell clustering, we integrated these data with our previously published lymphoma dataset that also included reactive lymphoid tissue controls^20^. After rigorous quality filtering and doublet removal, 185,280 cells were retained for downstream analysis. Following batch correction by harmony, cluster s were defined as malignant and non-malignant B-cells, T and NK cells, myeloid cells, and erythrocytes (Figure S1). The relative number of myeloid cells obtained was low, likely due to loss during physical disaggregation. Thus, we focused on the T-cell compartment, which consisted of 58.44% of the non-malignant cells within the rrLBCL tumors.

Re-clustering of proliferating T-cells identified CD4 and CD8 memory subsets and a CD8 effector subset (Figure 1A-C; Table S2). Within resting CD4 T-cells, we identified naïve (high *IL7R*, *SELL*, *LEF1*), ribosomal gene expression (RP_HI_), T follicular helper cells (T_FH_; high *CXCL13*, *CXCR5*, *PDCD1*), cytotoxic CD4 T-cells (CD4_Cyt_; high *GZMK*, *GZMA*, *PRF1* and cytotoxicity signature), and regulatory T-cells (T_Reg_; high *FOXP3*, *IL2RA*, *CTLA4*) (Figure 1A-C; Table S3). Within resting CD8 T-cells we observed CD8 effector (CD8_Eff_; high *GNLY*, *GZMH*, *KLRD1* and cytotoxicity signature), CD8 memory (CD8_Mem_; high memory signature, *IL7R*, *SELL*), KLRG1+ (high *KLRG1*, *GZMM*, *CXCR4*), terminally exhausted CD8 T-cells (CD8_TEX_; low T-cell dysfunction signature, high *TOX*, *TIGIT*, *PDCD1*), and dysfunctional CD8 T-cells (CD8_Dys_, high T-cell dysfunction signature, high *LAG3*, *HAVCR2*, *CD38*) (Figure 1A-C; Table S4). Although the CD8_TEX_ cluster and CD8_Dys_ cluster both express high levels of *TOX* and co-inhibitory molecules such as *TIGIT* and *PDCD1* (PD1), the CD8_Dys_ cluster additionally showed high expression of co-inhibitory molecules such as *CTLA4*, *LAG3,* and *HAVCR2* (TIM3) as well as other genes associated with T-cell dysfunction such as *CD38*, *PRDM1*, and *BATF*^21–23^. The CD8_Dys_ subset also likely represents a state of terminal exhaustion according to an assessment of previously defined marker genes (Figure S1)^24^. An exploratory analysis of compositional differences within the T-cell compartment between CAR T naïve and relapsed tumors showed significantly higher frequencies of CD8 _Dys_ (Fisher FDR=8.97×10^−34^), resting CD8_Eff_ (Fisher FDR=2.04×10^−9^), T_Reg_ (Fisher FDR=1.20×10^−5^) and CD4_Cyt_ cells (Fisher FDR=4.07×10^−5^), and significantly lower frequencies of resting CD8_Mem_ (Fisher FDR=1.4×10^−5^), KLRG1+, and naïve CD4 T-cells (Fisher FDR=8.38×10-219) as the most prevailing trends in CAR T relapsed tumors compared to CAR T naïve tumors (Figure S1; Table S5). Assessment of CAR expression in cells from CAR T relapsed tumors (Figure 1D-E) showed that the CAR+ cells were significantly enriched within the CD8_Dys_ cluster (520 of 695 CAR+ cells vs 866 of 1,690 CAR-cells: Fisher P < 0.0001) and the proliferating CD8 effector cluster (1015/2564 cells from CAR T relapsed tumors). Evaluation of TCR clonotypes identified a high degree of clonal expansion within the CD8_Dys_ and proliferating CD8 effector clusters, particularly within the CAR+ fraction of these clusters (Figure 1F-G). Furthermore, major hyperexpanded TCR clonotypes were present within the CAR+ fraction of both the proliferating CD8_Eff_ and resting CD8_Dys_ clusters (Figure 1H). Considering this, we performed trajectory analysis rooted in the resting CD8_Eff_ cluster, which showed a predicted temporal relationship terminating at the CD8_Dys_ cluster (Figure 1I). Together, these observations therefore indicate that ongoing target engagement and clonal expansion of CAR+ effector CD8 T-cells lead to T-cell dysfunction and the accumulation of clonally expanded CD8_Dys_ at the time of progression from CAR T-cell therapy.

**Figure 1:**
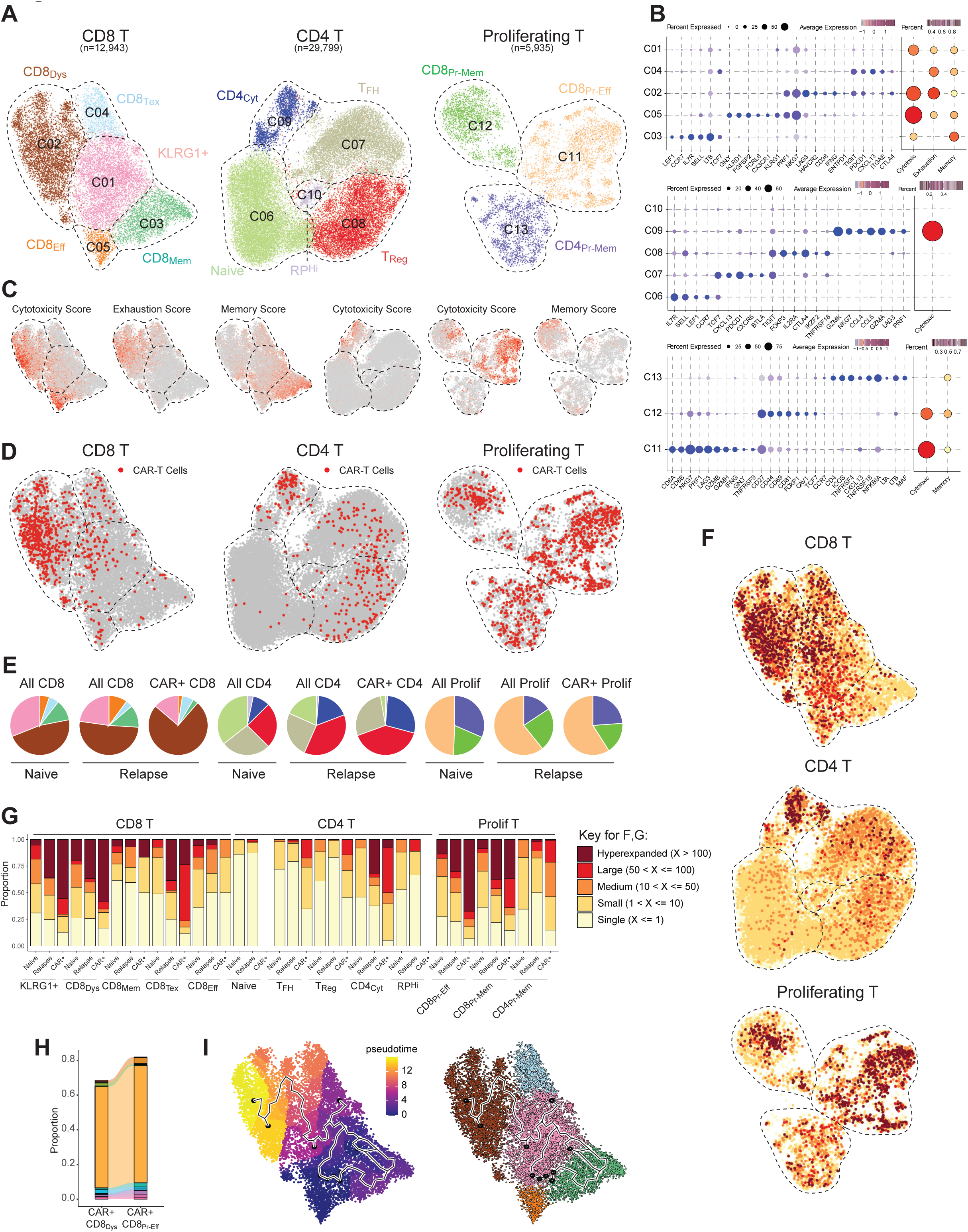
Clonally expanded dysfunctional CD8 T-cells are associated with CAR T-cell refractoriness in rrLBCL. **A**) UMAP projections of T-cell clusters (CD8, CD4, and proliferating) and corresponding T-cell subsets in the tumor microenvironment (TME) of relapsed/refractory large B cell lymphoma (rrLBCL) tumors from CAR T naïve patients (n=5) and those refractory to CAR T-cell therapy (n=7). DYS, dysfunctional; TEX, terminal exhaustion; FH, follicular helper; REG, regulatory. **B)** Gene expression bubble plot of T-cell subsets comprising the TME of rrLBCL tumors. Top, Middle, Bottom: Gene signature and subsets of CD8, CD4, and proliferating T-cells, respectively. **C)** Cytotoxic, exhaustion, and/or memory score expression projected onto T-cell cluster UMAPs from (A). **D)** CAR transcript expression in T-cells from TME projected onto T-cell cluster UMAPs from (A). **E)** Proportion of T-cell subsets within each group of T-cells in the TME of CAR T naïve patients (naïve) compared to those that relapsed following CAR T-cell therapy (relapsed). **F)** UMAP of TCR clonal expansion of T-cell clusters from (A). **G)** Bar chart illustrating TCR clonal expansion of T-cell subsets by CAR T naïve and relapsed patients and within CAR+ T-cells. **H)** Clonal tracking of top TCR clonotypes showing shared clonotypes between CD8_Dys_ and proliferating CD8 effector T-cells, respectively. **I.** Pseuodtime analysis demonstrating trajectory of CD8 T-cells terminating at CD8_Dys_ cluster. See also Figures S1A-C and Tables S1-S5

### Pegenzileukin rescues CD8 T-cell dysfunction driven by chronic antigen stimulation

Given the association between CD8 T-cell dysfunction and CAR T-cell relapse, we developed an *in vitro* model to explore CD8 T-cell dysfunction within a classical antigen/TCR-driven system using serial exposure of T-cells to HLA class I restricted peptides^25^. In brief, healthy donor CD8 T-cells were serially exposed to antigen on day 0, 4, 8 and 12 via dendritic cells primed with peptide, which generated dysfunctional T-cells as evidenced by a marked reduction in interferon gamma (IFNγ) production and cell proliferation compared with CD8 T-cells that were exposed to antigen only once (CD8+Pentamer+ IFNγ+: 15% vs 87%, Figure 2A). We therefore utilized this system to test whether pegenzileukin would be capable of rescuing (treatment post-exposure) or preventing (treatment concomitant with exposure) antigen-driven CD8 T-cell dysfunction.

**Figure 2:**
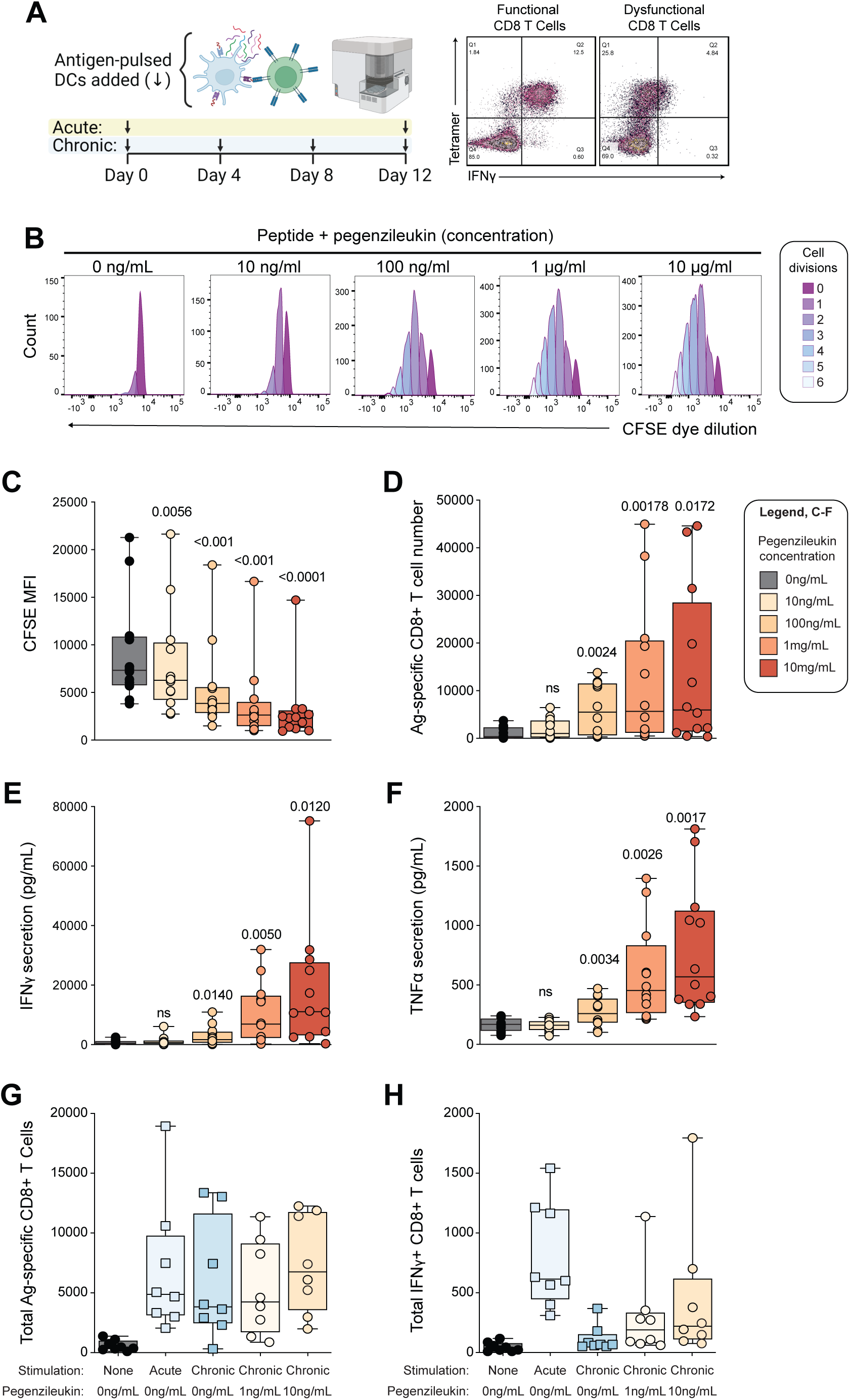
Pegenzileukin rescues CD8 T-cell dysfunction driven by chronic antigen stimulation. **A**) Schematic overview of chronic antigen stimulation assay and flow cytometry of interferon gamma (IFNγ) production from functional (left) and dysfunctional CD8+ T-cells (right) following HLA class I restricted peptide (antigen) exposure (one vs. four rounds of exposure, respectively). **B)** Flow cytometry analysis of antigen-specific CD8+ T-cell proliferation following administration of various doses of pegenzileukin to antigen-stimulated T-cells by CFSE dye dilution after multiple antigen stimulation. **C)** Boxplot of mean fluorometric intensity (MFI) of CFSE from antigen-stimulated CD8+ T-cells following treatment with various doses of pegenzileukin from (B). **D)** Boxplot of absolute number of antigen-stimulated CD8+ T-cells following treatment with various doses of pegenzileukin after multiple antigen stimulation. **E)** Boxplot of IFNγ production from antigen-stimulated T-cells treated with various doses of pegenzileukin after multiple antigen stimulation. **F)** Boxplot of MFI of tumor necrosis factor alpha (TNFα) production from antigen-stimulated T-cells treated with various doses of pegenzileukin after multiple antigen stimulation. **G)** Comparison of the absolute number of antigen specific CD8+ T-cells exposed to no, single, or multiple rounds of antigen and T-cells pre-treated with 10 ng/ml or 1 ng/ml dose of pegenzileukin then followed by multiple rounds of antigen exposure. **H)** Comparison of the absolute number of IFNγ producing CD8+ T-cells exposed to no, single, or multiple rounds of antigen and T-cells pre-treated with 10 ng/ml or 1 ng/ml dose of pegenzileukin followed by multiple rounds of antigen exposure. P-values shown are Bonferroni adjusted where adjusted p>0.05 = ns.

Rescue of CD8 T-cell dysfunction was evaluated in 12 independent healthy donors stimulated with antigen on days 0, 4 and 8, then stained with CFSE prior to the day 12 stimulation. Cells were then cultured in the presence or absence of pegenzileukin from days 12 to 19 and proliferation was measured by CFSE dye dilution. The addition of pegenzileukin at doses of 10μg/mL, 1μg/mL, 100ng/mL, and 10ng/ml significantly improved the proliferative capacity of CD8 T-cells in a dose-dependent manner with reference to untreated controls (Figure 2B-C). This was also confirmed using absolute cell numbers of antigen specific CD8 T-cells, which were significantly increased with the addition of pegenzileukin at all concentrations except 10 ng/ml (Figure 2D). Furthermore, the addition of pegenzileukin significantly increased the secretion of effector cytokines including IFNγ and TNFα (Figure 2E-F).

Prevention of T-cell dysfunction was evaluated by treatment of CD8 T-cells with pegenzileukin at doses of either 1 ng/mL or 10ng/mL during serial antigen stimulations over 12 days. We compared the pegenzileukin-treated to untreated cells exposed to multiple antigen stimulation (1-1-1), untreated cells exposed to a single antigen stimulation (1-0-0), or cells not exposed to antigen. The number of total antigen specific CD8 T-cells did not differ when comparing the single versus multiple antigen-stimulated T-cells and was unchanged with the addition of pegenzileukin (Figure 2G). However, the number of functional CD8 T-cells, as measured by IFNγ production, was significantly reduced by multiple antigen stimulations compared to the single antigen stimulation, which was mitigated by addition of 10ng/mL pegenzileukin (Figure 2H). Therefore, in the setting of chronic antigen stimulation, pegenzileukin can rescue or prevent CD8 T-cell dysfunction.

### LBCL patient-derived CAR T-cells are prone to T-cell dysfunction following chronic CAR-driven stimulation

Dysfunction of autologous CAR T-cells from rrLBCL patients is likely contributed to by both the intrinsic functional state of the T-cells harvested at the time of leukapheresis, resulting in variabilities within the functional state of CAR T-cell infusion products, and by resistance of tumor cells to killing which has been shown to promote chronic stimulation and exhaustion^26^. We therefore developed an *in vitro* model of CAR T-cell dysfunction using (i) T-cells from rrLBCL patients taken at the time of leukapheresis for standard of care CAR T-cell therapy, and (ii) utilizing the RL (CD19+ LBCL) cell line as targets due to its resistance to T-cell killing^27^. Specifically, RL target-cells were co-cultured with CD19 CAR T-cells at an effector to target ratio of 1:1 at the start of the assay, and an equivalent number of fresh target-cells were added every 2 days thereafter over a period of 8 days (Figure 3A). We used this assay to evaluate whether CD19 CAR T-cells from rrLBCL patients (n=3), taken at the time of leukapheresis for standard of care CAR T-cell therapy, were more prone to T-cell dysfunction compared to those generated from healthy donors (n=3, age 60+ donors). This was evaluated through a 22-marker cell surface immunophenotyping performed at baseline, day 4, and day 8, and measurement of effector function (target cell killing) and T-cell expansion every 2 days by counting T-cells and tumor cells via flow cytometry.

**Figure 3.**
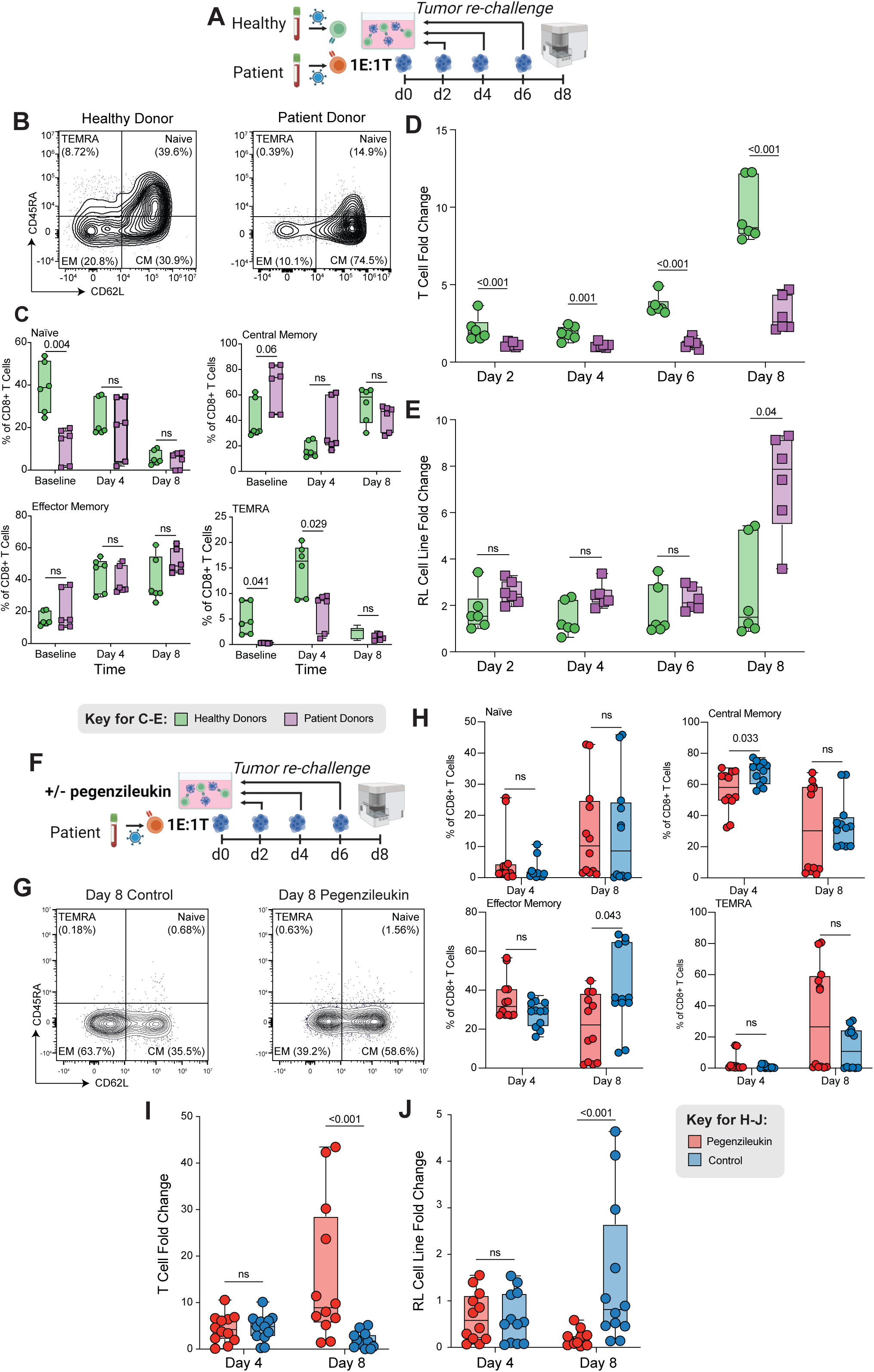
LBCL patient-derived CAR T-cells are prone to T-cell dysfunction following chronic CAR-driven stimulation and rescued by pegenzileukin. **A**) Schematic of CAR T-cell tumor re-challenge assay with healthy donor compared with patient derived CAR T-cells. **B)** Flow cytometry analysis of the proportion of CD8 T-cell differentiation states at baseline prior to tumor rechallenge assay. Naïve (CD45RA+CD62L+), effector memory expressing CD45RA (TEMRA) (CD45RA+CD62L-), central memory (CD45RA-CD62L+), and effector memory (CD45RA-CD62L). TEMRA, terminally exhausted memory cells re-expressing CD45RA. **C)** Boxplot from flow analysis comparing change in the proportion of various differentiation states between healthy donor (green) and patient derived (purple) CD8+ T-cells from baseline to day 8 in tumor re-challenge assay. **D)** Expansion from baseline of T-cells derived from healthy donors (green) compared to those derived from patients with rrLBCL (purple) following repeated RL tumor exposure in a tumor re-challenge assay. N=3 healthy and patient CAR T donors, respectively. **E)** Expansion of RL tumor cells from baseline *in vitro* in co-culture with healthy donor derived T-cells (green) compared to patient derived T-cells (purple). **F)** Schematic of CAR T-cell tumor re-challenge assay with patient derived CAR T-cells with or without the addition of pegenzileukin. **G)** Flow cytometry analysis comparing the proportion of patient donor T-cell differentiation states between the control (top) and pegenzileukin treated (bottom) groups at day 8 of serial tumor re-challenge assay. **H)** Boxplot from flow analysis comparing changes in the proportion of various CD8+ T-cell differentiation states between control (blue) and pegenzileukin treated (red) groups at days 4 and 8 in the tumor re-challenge assay. **I)** Expansion from baseline of patient derived T-cells treated with pegenzileukin (red) or control (blue) in an 8-day serial tumor re-challenge assay. N=6 R/R lymphoma patient CAR T donors. **J)** Expansion from baseline of RL tumor in co-culture with pegenzileukin treated patient derived CAR T-cells (red) compared to non-treated CAR T-cells (blue) in an 8-day serial tumor re-challenge assay. See also Figures S2 and S3 P-values shown are Bonferroni adjusted where adjusted p>0.05 = ns.

Healthy donor T-cells were highlighted by distinct differences in CD8 memory compartments, including more naïve (CD45RA+CD62L+) and effector memory expressing CD45RA (TEMRA) (CD45RA+CD62L-) CD8 T-cells and less central memory (CD45RA-CD62L+) CD8 T-cells, present prior to exposure of CAR T-cells to target cells (Figure 3B-C). However, these compartments were not significantly different following challenge with target cells (Figure 3C). A similar trend was observed in CD4 T-cells with more naïve and TEMRA CD4 T-cells, and less central memory CD4 T-cells (Figure S2A). Although we observed slightly elevated TIGIT expression in patient derived CD8+ effector memory T-cells on day 4 and day 8, and induction of increased expression of OX40 on CD8+ effector memory T-cells from healthy donor CAR T-cells (Figure S2B), the serial rechallenge of CAR T-cells was not accompanied by significant immunophenotypic differences in either the CD8 or CD4 compartment between rrLBCL patient donors and healthy donors (Figure S2B-C). Despite this, we noted marked functional differences between rrLBCL patient donor and healthy donor CAR T-cells. At each time point in the assay, healthy donor T-cells exhibited increased expansion with maximal (9.7-fold) expansion reached by day 8, in contrast to minimal expansion (3.1-fold) for patient-derived CAR T-cells (q=0.0007, Figure 3D). Furthermore, the serial re-challenge rrLBCL patient-derived CAR T-cells drove a collapse in effector function at day 8 as evidenced by failure to control tumor cell outgrowth (7.3-fold increase in RL cells over baseline), which was not observed with healthy donor-derived CAR T-cells (2.6-fold increase in RL cells over baseline) (q=0.04, Figure 3E). Thus, CAR T-cells derived from rrLBCL patient PBMCs taken at the time of apheresis for CAR T-cell therapy are prone to dysfunction but do not exhibit measurable changes in cell surface protein expression of therapeutically targetable co-inhibitory molecules, suggesting that alternative strategies for rescuing CAR T-cell function may be needed.

### Rescue of CAR T-cell dysfunction *in vitro* by pegenzileukin

We next tested whether pegenzileukin may be capable of rescuing the function of patient-derived CAR T-cells in our *in vitro* assay. In short-term (24 and 48h) assays with a single challenge of tumor cells at a 1:1 effector to target ratio, pegenzileukin did not increase cell killing at either 24 (Figure S3A) or 48h (Figure S3B) but did increase T-cell proliferation at both 24 (Figure S3C) and 48h time point (Figure S3D). In an 8-day serial re-challenge with 6 independent rrLBCL donors (Figure 3F), the scenario in which we observed the greatest degree of T-cell dysfunction, we evaluated both immunophenotypic and functional changes associated with pegenzileukin treatment. Continuous high-level stimulation by IL-2 is capable of inducing exhaustion of tumor-reactive T-cells, including the up-regulation of co-inhibitory molecules such as PD-1 and LAG-3^28^. We observed a modest reduction in the proportion of central memory CD8 T-cells on day 4, and a modest decrease in effector memory CD8 T-cells on day 8 in pegenzileukin treated samples compared to controls (Figure 3G-H). However, there was a large degree of patient-to-patient heterogeneity in these phenotypes, likely linked with the heterogeneous treatment histories and disease biology between patients. No significant differences were observed for CD4 T-cell populations at either day 4 or 8 (Figure S3E). Detailed immunophenotypic characterization of T-cells at the mid-point (day 4) and endpoint (day 8) of the assay showed only minor differences in the composition and expression of co-inhibitory or co-stimulatory markers between treated and untreated groups in both the CD8 (Figure S3F) and CD4 compartments (Figure S3G). However, the addition of pegenzileukin resulted in significantly increased CAR T-cell expansion by day 8 (average=15.9-fold vs 2.0-fold in untreated controls; p=0.0004) (Figure 3I). In addition, pegenzileukin allowed continued lymphoma cell killing by day 8, whereas CAR T-cells not exposed to pegenzileukin lost the ability to control tumor cell outgrowth (p=0.0003; Figure 3J). Pegenzileukin is therefore capable of preventing T-cell dysfunction in patient-derived CAR T-cells following serial tumor re-challenge without significant effects on T-cell immunophenotype.

### Pegenzileukin leads to *in vivo* expansion of CAR T-cells and sustained tumor control

We next tested the activity of pegenzileukin within an *in vivo* lymphoma model in which CAR T-cells provide some control of the tumor but with incomplete clearance. Luciferized Raji cells were injected on day 0 and mice were treated with either untransduced T-cells or CD19/CD22 CAR T-cells on day 10 with the addition of 3 weekly doses of pegenzileukin at a dose of 6 mg/kg or control starting on day 9. Untreated mice (tumor only), mice treated with untransduced T-cells without pegenzileukin, and mice treated with untransduced T-cells in combination with pegenzileukin had similar tumor growth kinetics with no effect of untransduced T-cells in either condition (Figure 4A). Treatment with CAR T-cells in the absence of pegenzileukin provided a measurable response marked by a plateau in tumor growth, but without a marked decline in tumor burden. In contrast, mice treated with CAR T in combination with pegenzileukin showed significantly improved tumor control by day 29 as measured by bioluminescence (p=0.047, Figure 4A-B). This was also supported by postmortem analysis of tumor cells within the lungs (Figure 4C and 4E) and liver (Figure 4D and 4F) of these mice using immunohistochemical staining for human CD20 (hCD20). Analysis of peripheral blood T-cells by flow cytometry also revealed a significant increase in the expansion of CD4+, CD8+, CD4+CAR+, and CD8+CAR+ T-cells in the peripheral blood of mice at days 21 and 28 in pegenzileukin treated mice compared to untreated mice (Figure 4G). Therefore, pegenzileukin significantly improves expansion and function of CAR T-cells in an *in vivo* model of sub-optimal tumor control.

**Figure 4.**
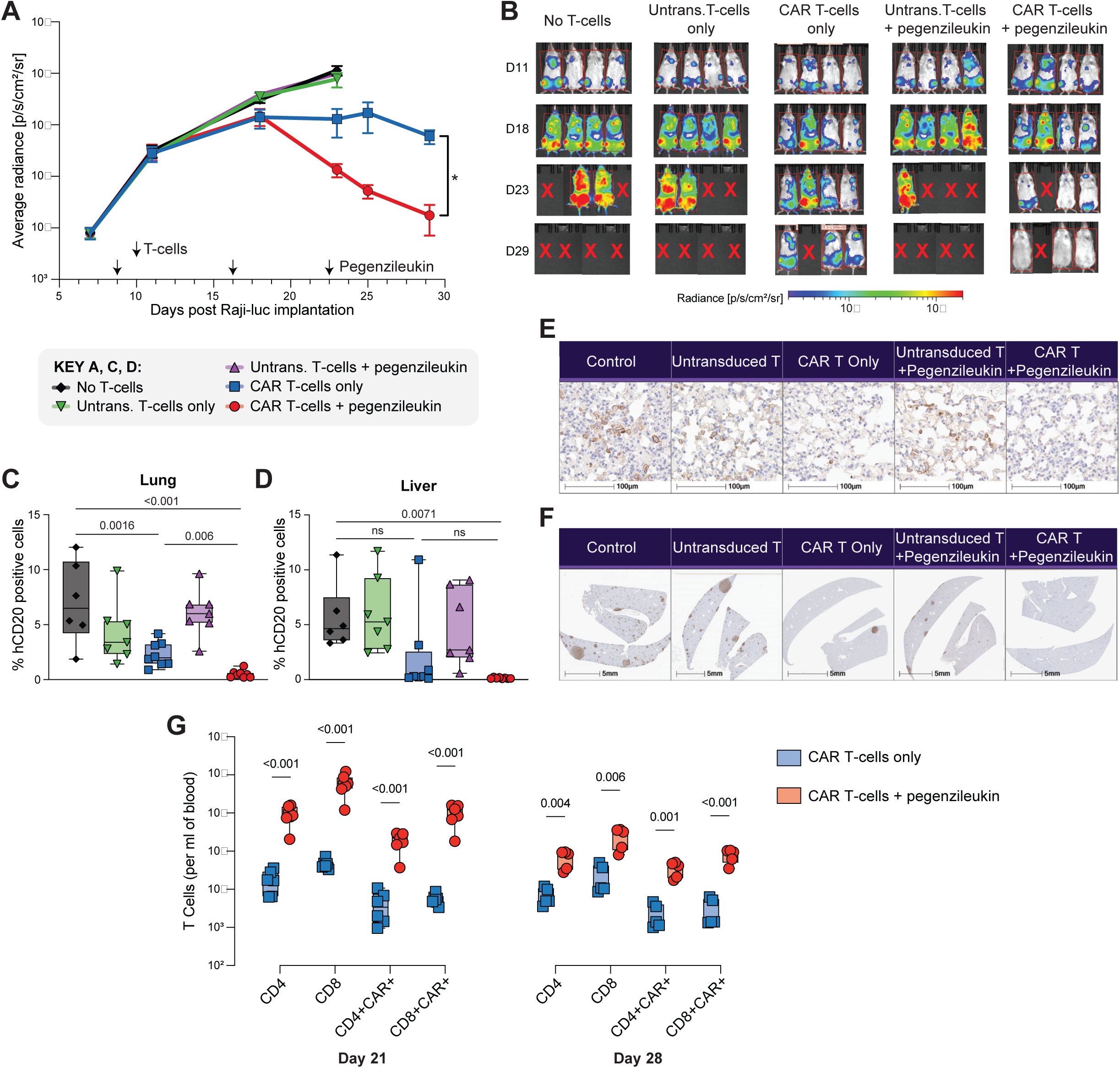
Pegenzileukin leads to *in vivo* expansion of CAR T-cells and sustained tumor control. **A**) Growth of luciferized Raji (Raji-luc) tumor cells as measured by bioluminescent imaging of mice treated with either untransduced T-cells, anti-CD19/CD22 CAR T-cells, untransduced T-cells with pegenzileukin, CAR T-cells with pegenzileukin, or vehicle only. Raji-luc given IV on day 0, T-cells given IV on day 10, and pegenzileukin given IV for 3 weekly doses starting on day 9. N=4 mice per group. **B)** Bioluminescence imaging of *in vivo* Raji-luc mice models with various treatment conditions. **C)** Bar graph reflecting percentage of human CD20 (hCD20) in lung tissue of *in vivo* Raji-luc mice models cells as a surrogate for tumor persistence. **D)** Bar graph reflecting percentage of hCD20 in liver tissue of in vivo Raji-luc mice models cells as a surrogate for tumor persistence. **E)** Postmortem immunohistochemical (IHC) staining of tumor cells in lung tissue from *in vivo* Raji-luc mice models from each treatment condition corresponding to (C). **F)** Postmortem IHC staining of tumor cells in liver tissue from *in vivo* Raji-luc mice models from each treatment condition corresponding to (D). **G)** Peripheral blood analysis from *in vivo* Raji-luc mice models comparing expansion of T-cells between pegenzileukin (red) and untreated groups (blue) at days 21 and 28 of experiment. P-values shown are Bonferroni adjusted where adjusted p>0.05 = ns.

## DISCUSSION

CAR T-cell failure is likely contributed to by many factors that are inextricably linked, including the quality of T-cells at the time leukapheresis, the subsequent quality of the CAR T-cell infusion product, tumor burden at the time of infusion, tumor immune microenvironment characteristics, and tumor-intrinsic mechanisms of resistance or escape from CAR T-cell killing or recognition. T-cell exhaustion/dysfunction has recurrently emerged as an important feature associated with CAR T-cell failure^6,29,30^ but has not been explored within tumors at the time of relapse following CAR T-cell therapy. Our scRNA-seq showed significant compositional differences between rrLBCL tumors that were CAR T naïve compared to those from post-CAR T relapse, including a marked clonal expansion and over-representation of dysfunctional CD8 T-cells that was particularly notable within the CAR+ compartment. Our ability to track CAR+ cells within these data, and to specifically track expanded TCR clonotypes of CAR+ cells between clusters, lead us to hypothesize that the clonal expansion and high rates of CD8 T-cell dysfunction enriched in post-CAR T-cell relapse tumors are likely actively acquired post-infusion in a process of tumor engagement and chronic stimulation, rather than being pre-existent. Further analysis of large cohorts of matched pre– and post-treatment samples are required to definitively prove this hypothesis. Furthermore, other features of the microenvironment such as increased frequencies of myeloid cells and T_Reg_ cells have been observed by others to be associated with poor outcome following CAR T-cell therapy^17,18,31,32^ and are also likely to play an important role in CAR T-cell failure.

The susceptibility of T-cells to terminal exhaustion or dysfunction is likely to be patient and disease specific. Both chronic antigen stimulation and replicative dysregulation of CAR T-cells leads to T-cell dysfunction associated with a loss of proliferation, terminal differentiation of effector T-cells, and resulting loss of anti-tumor cytotoxicity^33,34^. In line with this, CAR T-cell infusion products with limited proliferative potential are associated with worse outcomes whereas less differentiated T-cells with superior proliferative capacity demonstrate better responses^30,35^. Here we show that lymphoma patient T-cells are significantly more prone to dysfunction than T-cells from healthy patients, corroborating recent findings comparing acute lymphoblastic leukemia and chronic lymphocytic leukemia patient samples to healthy donors^36^. We therefore utilized patient derived CAR T-cells in an *in vitro* model system of serial re-challenge with T-cell-resistant target cells to test the efficacy of pegenzileukin, a non-alpha IL-2R agonist, which we hypothesized would promote CAR T-cell expansion and prevent the terminal dysfunction/exhaustion. Within these assays, pegenzileukin was able to restore the proliferative potential of T-cells and promote tumor control. This was supported within orthogonal functional models of *in vitro* antigen:TCR-driven assays of T-cell dysfunction, and an *in vivo* lymphoma model of sub-optimal CAR T-cell activity. This is consistent with prior results in murine models of melanoma in which pegenzileukin led to an increase in the proliferation of, and enhanced anti-tumor cytotoxicity of NK and CD8 T-cells through activation of STAT5^19^. Taken together, these data highlight pegenzileukin and other similar interventions as possible therapeutic approaches for improving CAR T-cell function and patient response. Pegenzileukin has shown to be well tolerated as a single agent and in combination with PD-1 blockade^37^. However, interventions such as these would likely be best suited for patients at a high risk of CAR T-cell failure, as determined by pre-treatment biomarkers^38–43^.

In summary, we have shown in a discovery analysis using single cell RNA-sequencing of primary CAR T relapse tumors, together with orthogonal functional validation using patient-derived CAR T-cells, that CD8 T-cell dysfunction is a major contributor to CAR T-cell failure in rrLBCL. Pegenzileukin, a non-alpha IL-2R agonist, was capable of restoring T-cell expansion and tumor control in the setting of both antigen-driven and CAR-driven T-cell dysfunction *in vitro* and *in vivo*. This highlights pegenzileukin and other therapeutics with similar mechanisms of action as attractive candidates for combination with CAR T-cell therapy in patients at a high risk of failure.

## ACKNOWLEDGEMENTS

This study was funded by CPRIT IIRA RP200385 and by Sanofi. PKR is supported by the Lymphoma Research Foundation Postdoctoral Fellowship Grant, the Lymphoma Research Foundation Lymphoma Scientific Research Mentoring Program, and a Conquer Cancer Foundation Young Investigator Award. INS is supported by a Conquer Cancer Foundation Young Investigator Award. MRG is supported by a Scholar award from the Leukemia and Lymphoma Society. The MD Anderson Lymphoma Tissue Bank is supported by KW Cares.

## STAR METHODS

### Patient samples and single cell RNA-sequencing

rrLBCL tumor specimens were obtained with informed consent in accordance with protocols approved by review board of University of Texas MD Anderson Cancer Center. All samples were mechanically dissociated into single-cell suspension. PBS containing 0.04% UltraPureTM Bovine Serum Albumin (BSA, 50mg/ml) was used for cell washing and resuspension to minimize cell losses and aggregation. Cell viability was assessed by trypan blue examination, and samples with more than 80% viable cells were chose for Chromium Single Cell Immune Profiling Solution, according to Chromium Single Cell 5’ Library & Gel Bead Kits User Guide (v1 Chemistry). Briefly, the final cell concentration was adjusted to ∼1000 cells/µl, single cell suspension mixed with reverse transcription (RT) master mixture were loaded on a 10X Genomics single cell instrument. GEMs were broken and single-strand cDNA was cleaned up with DynaBeads. Amplified cDNA quality and quantity were assessed by High Sensitivity D5000 DNA Screen Tape analysis (Agilent Technologies) and Qubit dsDNA HS Assay Kit (Thermo Fisher Scientific).

#### Single-cell RNA-Library construction and sequencing

We used the 10X Genomics Single-Cell 5’ Library Kit (PN-1000020) to construct indexed sequencing libraries following the manufacturer’s protocol. Sequencing with indexing the Chromium i7 Samplex index Kit (PN-120262) was conducted on an Illumina Hiseq4000 sequencer with 2×100 bp paired reads to achieve a depth of at least 50,000 read pairs per cell.

#### Single-cell V(D)J Enrichment library construction and sequencing

Chromium Single Cell V(D)J Enrichment Kit (Human B-cell, PN-1000016; Human T-cell, PN-1000005) from 10X Genomics was employed to enrich immune repertoire, T-cell receptor (TCR) or B-cell immunoglobulin (Ig) transcripts. 50 ng of enriched BCR or TCR products was used for library construction, according to manufacturer’s protocol. Single Cell V(D)J enriched library was indexed by Chromium i7 Samplex index Kit (PN-120262) and multiple samples were pooled in a sequence lane at 150 x 150 bp with minimum 5,000 read pairs per cell.

### Single-cell RNA-sequencing analysis

#### Data processing and quality control

The raw scRNA-seq data underwent pre-processing, including demultiplexing cellular barcodes, read alignment, and gene count matrix generation, using Cell Ranger Single Cell Software Suite (version 3.1.0) from 10× Genomics. Quality control (QC) metrics were meticulously evaluated to ensure high-quality data for downstream analyses. To achieve this, cells with low complexity libraries (detecting transcripts aligned to < 200 genes) were filtered out for basic quality filtering. This filter helps to remove cell debris, empty drops, and low-quality cells. Additionally, cells likely to be dying or apoptotic were excluded, where > 15% of transcripts originated from the mitochondrial genome.

#### Doublet detection and removal

Doublets or multiplets that were likely present in the scRNA-seq data were detected and carefully removed through a multi-step approach as described in recent studies. In brief, doublets or multiplets were identified through the following methods: 1) library complexity, where cells with high-complexity libraries detecting transcripts aligned to > 6500 genes (the top ∼1% outliers in the distribution of genes detected per cell) were removed. 2) Cluster distribution and marker gene expression, where some doublets or multiplets can form distinct clusters with hybrid expression features, exhibiting an aberrantly high gene count. Expression levels and proportions of canonical lineage-related marker genes in each identified cluster were reviewed, and clusters co-expressing discrepant lineage markers (e.g., B-cell clusters showed expression of T, myeloid, or stromal cell markers) were removed. 3) Integration of scTCR/BCR-seq data, where cells with both productive T and B cell receptors (TCR/BCR) or had ≥ 2 productive TCRs/BCRs were removed. Cells of non-T/B cell clusters with productive TCRs or BCRs were also excluded. 4) Cells co-expressing ≥ 2 distinct lineage markers, where some doublets or multiplets did not form separate cell clusters but were dispersed across the UMAP plots. To identify and clean doublets missed in the previous steps, canonical marker gene expression in defined cell clusters were inspected, and cells co-expressing discrepant lineage-specific markers were removed (e.g., cells in the T-cell cluster showed expression of both CD4 and CD8A/B). Steps 2) and 4) were repeated multiple times to filter out most barcodes associated with cell doublets. After doublets removal, a total of 185,280 cells were retained for downstream analyses. Normalization of library size was performed in Seurat on the filtered gene-cell matrix to obtain normalized UMI counts.

#### Batch effect evaluation and correction

To assess possible batch effects, we performed a statistical analysis using *k-BET*, a robust and sensitive k-nearest neighbor batch-effect test R package. Major non-malignant cell types from all samples were analyzed separately with default parameters, as described in our previous studies. Subsequently, we utilized *Harmony*, one of the top-ranked methods for batch effect correction, to iteratively remove the identified batch effects present in the principal component analysis (PCA) space. During the clustering of major TME cell lineages, Harmony was executed with default parameters. Finally, we conducted an assessment to evaluate Harmony’s performance in terms of mixing batches while preserving cell-type purity.

#### Unsupervised clustering and sub-clustering analysis

To identify highly variable genes (HVGs) for unsupervised cell clustering, we applied Seurat (version 4.0.0) to the normalized gene-cell matrix. We performed principal component analysis (PCA) on the top 2,000 HVGs and generated an elbow plot using Seurat’s *ElbowPlot* function to determine the number of significant principal components (PCs) required. The *FindNeighbors* function of Seurat was used to construct the Shared Nearest Neighbor (SNN) Graph based on unsupervised clustering performed with Seurat’s *FindClusters* function. We conducted multiple rounds of clustering and sub-clustering analysis to identify major cell lineages and distinguish distinct cell transcriptional states within each major cell type. We used Uniform Manifold Approximation and Projection (UMAP) with Seurat’s *RunUMAP* function to perform dimensionality reduction and 2-D visualization of the single-cell clusters, using the same number of PCs used for clustering. We checked for each cell type whether additional cell subclusters represented the same cell state without showing any unique features to determine whether cells were over-clustered. We tested different resolution and PC parameters for unsupervised clustering, reviewed the resulting UMAP plots and cluster signature genes, and used these results to determine the optimal number of clusters and guide the proper clustering of our scRNA-seq data.

#### Determination of cell types and states

The cell types and states were determined based on the *Harmony*-defined cell clusters. To identify major cell types and subpopulations, two rounds of clustering (clustering and sub-clustering) were performed on all cells, focusing on CD4 T, CD8 T, proliferating T, NK, B/Plasma, myeloid, and malignant cells. Both rounds of clustering relied on determining the 50-nearest neighbors of each cell based on 30 PCs to construct SNN graphs. Then, using the *FindMarkers* function in Seurat, differentially expressed genes (DEGs) for each cluster were identified and filtered based on several criteria: expression in at least 25% of cells in the more abundant group, a fold change in expression of at least 1.2, and FDR adjusted p-values (*p.adj*) less than or equal to 0.05. Additionally, mitochondrial and ribosomal genes were excluded from the DEG lists. Bubble plots were generated for selected DEGs and a suggested set of canonical markers for various kinds of cell lineages. Multiple layers of information, including cluster distribution, cluster-specific gene expression (particularly the top 50 DEGs), canonical cell lineage markers, and functional gene signatures, were integrated and carefully reviewed to define cell types and transcriptomic states. This process was described in our previous report. Finally, platelets and red blood cells were excluded from downstream analyses.

#### Identification of CAR-positive T-cells

Gene dropouts are a common occurrence in scRNA-seq data, which poses a challenge to identifying cell types. To enhance the accuracy of identifying CAR-positive T-cells, we utilized CapID sequencing data to distinguish positive and negative expression. In addition, we used a novel GRCh38 assembly (hg38_gencode34_CAR5p_v1), specifically designed for identifying CAR-positive cells, as a reference. By analyzing both scRNA-seq and CapID sequencing data, we were able to identify CAR-positive T-cells by detecting the presence of CAR-specific sequence contigs (Yescarta and FMC63-CD19scFV) in the aligned reads. This approach successfully defined a total of 2,305 QC-passed T-cells as CAR positive.

#### Scoring of curated gene signatures

To infer the functional states of T-cell subsets, we collected a list of curated gene signatures from published studies, including the Memory, Cytotoxic, and Exhaustion signatures. The signature scores for each cell were computed using the R package *AUCell* in a manner as described previously.

#### scTCR-seq data assembly, paired clonotype calling, TCR clonality analysis, and integration with T-cell phenotypes

The TCR reconstruction and paired TCR clonotype calling for the V(D)J pipeline were performed using Cell Ranger v3.1.0. The GRCh38 assembly in Ensembl (refdata-cellranger-vdj-GRCh38-alts-ensembl-2.0.0) was used as the reference. CDR3 motifs were identified, and only productive V(D)J rearrangements were analyzed. The clonal fraction of each identified clonotype was calculated to quantify the degree of T-cell clonal expansion. The T-cell clonotype landscape was assessed using the identified clonotypes. The clonality and diversity of T-cells were measured using the Simpson Clonality and Shannon Entropy indices, respectively, using the *scRepertoire* R package. The TCR clonotype data were integrated with T-cell transcriptional profiles at the single-cell level based on their shared unique cell barcodes.

#### Statistical analyses

In addition to the algorithms and statistical analyses mentioned above, we conducted all other basic statistical analyses in the R statistical environment (version 4.0.0). All statistical tests performed in this study were two-sided. Fisher’s exact test was employed to compare the frequencies of cell subsets between treatment-naïve and refractory samples. To account for multiple hypothesis testing, we applied the Benjamini-Hochberg method to correct *p*-values, and the false discovery rates (FDR *q*-values) were calculated. We considered results to be statistically significant if the *p*-value or FDR *q*-value was < 0.05. When the p-value reported by R (version 4.0.0) was smaller than 2e-16, it was reported as “*p* < 2×10^−16^”.

### In vitro Assays

#### Cell lines and Culture Conditions

All tumor cell lines (RL) used in this study were purchased from ATCC and validated to be free from mycoplasma from PCR. The tumor cell lines were cultured in RPMI 1640 (Corning). The RPMI-1640 media was supplemented with 10% heat-inactivated FBS (Gibco).

#### Patients and Specimens

Human peripheral blood mononuclear cells (PBMCs) from healthy and patient donors were obtained from the University of Texas MD Anderson Cancer Center using an Institutional Review Board (IRB) approved protocol.

#### Construction of Chimeric Antigen Receptor (CAR)

The structure of CD19 CAR was incorporated with the scFv derived from the FMC63 antibody, as well as the CD28 and CD3ζ signaling domains.

#### Lentiviral Vector Production and Transduction

Lentiviral vectors containing a transgene encoding CD19 CAR were produced. Lenti-293T cell lines were cultured using DMEM (Corning) supplemented with 10% FBS until 90% confluency. The encoding lentiviral supernatants for CD19 CAR were produced via transfection of the Lenti-293T cell line using LipofectamineTM 3000 (ThermoFisher Scientific) with the plasmids encoding the CARs and the Lentivirus envelope protein. Supernatants were collected 48 and 72 hours post transfection and viral confirmation and quantification done using Lenti-X GoStix Plus (Takara Bio).

PBMCs were depleted of monocytes using ImunoCult CD3/CD28 T-cell activator and 200 IU/mL IL-2 (Genscript) for 72 hours at a concentration of 1×10^6^ cells/ml of CAR T-cell media. Activated T-cells were then lentivirus transduced with the addition of Vectofusion-1 (Miltenyi Biotec) and cultured in Opti-MEM (Gibco) for 12 hours. Media was then changed to CAR T-cell media with 200 IU/mL human IL-2 (Genscript). Medium and IL-2 was replaced every 2 days. Transduction efficiencies were tested 72 hours after transduction and were routinely ∼50-70% for CARs. CAR T-cells were all used in in vitro assays days 6-8 post T-cell activation.

#### CD8+ T-cell stimulation and development of exhausted T-cells

CD8+ T-cells were isolated from frozen PBMCs by negative magnetic bead selection according to the manufacturer’s instructions (STEMCELL Technologies). CD8+ T-cells were co-cultured with autologous DCs previously primed with antiviral HLA-restricted peptides (BioSynthesis, Inc. Lewisville, TX, USA) at a ratio of 60:1 and incubated at 37oC and 5% CO2 in X-VIVO 15 serum-free media (Fisher Scientific) for 12-14 days. Exhausted CD8+ T-cells were developed by additional rounds of peptide stimulation during the assay. CD8+ T-cells that received a single peptide stimulation (functional) or multiple, repeated peptide stimulations (exhausted) were harvested, mixed with fresh peptide-pulsed autologous DC for 5 hours in the presence of Brefeldin A (Sigma-Aldrich) and stained with HLA Class I-restricted pentamers (Proimmune, Sarasota, FL, USA), phenotypic and functional markers of T-cell exhaustion via flow cytometry (manuscript in preparation).

#### Evaluation of Pegenzileuken in exhausted CD8 T-cells

Exhausted CD8 T-cells from 12-day co-cultures were harvested, labelled with Carboxyfluorescein diacetate succinimidyl ester (CFSE) (Life Technologies) using established protocols and re-stimulated with fresh autologous APCs pre-primed with the appropriate HLA Class I-restricted peptide and treated with various doses of pegenzileukin (Sanofi), recombinant human cytokine, or peptide alone (antigen only) at 37°C in 5% CO2 for an additional 7 days. Proliferation was assessed by flow cytometry evaluating the number of HLA-pentamer+ Ag-specific CD8 T-cells and the extent of CFSE dye dilution. Phenotypic markers of T-cell exhaustion were evaluated on Ag-specific CD8 T-cells marked by HLA Class I-restricted pentamers. Supernatants from the various 7-day re-stimulation CD8 T-cell culture conditions were analyzed for cytokine expression using the Milliplex® Human Cytokine Custom 17 Plex detection system (EMD Millipore) (Sigma-Aldrich), per the manufacturer’s protocol, and cytokine responses were measured on BioPlex™/ Luminex Systems and data was acquisitioned in Bio-Plex® Manager Software (Bio-Rad) (*manuscript in preparation*).

#### Tumor Re-Challenge Assays

Tumor cells (RL) and CAR T-cells (healthy or patient donors) were co-cultured at an effector to tumor ratio (E:T) of 1:1 7-10 days following initial activation of T-cells (4-7 days following CAR transduction). Duplicate conditions were performed, and cells were cultured in RPMI 1640 (Corning) supplemented with 10% FBS in 6 well culture plates. The co-culture assays were placed in incubation for 48 hours at 37°C in 5% CO2. Every 48 hours, each sample was centrifuged for the removal of debris, fresh media was added, and fresh tumor cells were replaced to co-culture prior to performing flow cytometry analysis. The assays were completed on day 8.

In tumor re-challenge assays using pegenzileukin (Sanofi), tumor cells and patient donor CAR T-cells were co-cultured at an effector to tumor ratio (E:T) of 1:1 7-10 days following initial thawing of cells (4-7 days following CAR transduction). Pegenzileukin (Sanofi) was added at baseline at a concentration of 0.4 μg/ml. Duplicate conditions were performed, and cells were cultured in RPMI 1640 (Corning) supplemented with 10% FBS in 6 well cell culture plates. The co-culture assays were placed in incubation for 48 hours at 37°C in 5% CO2. Every 48 hours, each sample was centrifuged for the removal of debris, fresh media and fresh tumor cells were replaced in co-culture and pegenzileukin (Sanofi) was added. The assays were completed on day 8.

#### Flow Cytometry

Cells were harvested at baseline for flow cytometry analysis prior to addition into co-culture and at days 4 and 8 of the re-challenge assay. Samples were stained separately for extracellular marker phenotyping using co-stimulatory and co-inhibitory panels. Human TruStain FcX™ (Biolegend) was utilized for Fc receptor blocking prior to the addition of extracellular antibodies. Antibody master mix was made the day of staining in BD Brilliant Stain Buffer (BD Biosciences). All antibodies were individually titrated for optimal staining and optimized antibody concentrations were used for cell staining. All cell samples were washed with FACS Buffer comprised of 1% bovine serum albumin (BSA) (Fisher Bioreagents) in PBS (Gibco). The samples were recorded using a Cytek Aurora and data analysis was performed on FlowJo v10.9 software.

### In Vivo Studies

#### Animals

All animal procedures were performed in accordance with the guidelines of the Animal Care and Use Committee of Sanofi. Following drug treatment, mice were observed daily for overall health including general appearance of the fur, mobility, and body weight. The animals were euthanized before they exhibited clinical signs and symptoms to avoid unnecessary pain and discomfort, according to internal ethical animal guidelines.

#### CAR T-cell generation

PBMCs (STEMCELL Technologies) were sent to Promab Biotechnologies for customized CAR T-cells production. CAR T-cells targeting CD19 and CD22 antigens were generated (ProMab Biotechnologies) with CD19scFV-CD22scFV-4-1BB-CD3zeta CAR construct. T-cells were isolated from PBMCs and activated with IL-2 and CD3/CD28 MacrobeadsTM (ProMab Biotechnologies) and transduced with lentivirus 24 hours and 48 hours following activation. Cells were passaged at 2–3 day intervals to maintain proper cell density. Starting on day 8, CAR expression was detected using flow cytometry (https://www.promab.com/car-t-cell-production). Isolated T-cells without lentivirus CAR transduction (untransduced) were used as control for CAR T *in vivo* studies.

#### Raji-Luc tumor model and In vivo drug treatment

Raji cell lines engineered to constitutively express Luciferase (Raji-Luc, Cellomics) were cultured in Dulbecco’s modified Eagle’s medium (DMEM) (Gibco) supplemented with 10% fetal bovine serum (Gibco) and Puromycin (0.5μg/mL, Gibco). Mice aged 7–8-week-old, female, NOD.Cg-Prkdcscid H2-K1tm1Bpe H2-Ab1em1Mvw H2-D1tm1Bpe Il2rgtm1Wjl/SzJ (NSG-MHC-DKO, Jackson Laboratories) were implanted with 0.5 × 106 Raji-Luc cells intravenously through tail vein injection. Tumor progression of Raji-Luc was monitored at least twice per week by in vivo bioluminescence imaging (BLI) using the Xenogen IVIS Spectrum Imaging System (Perkin Elmer). Mice were randomized and assigned to treatment groups according to baseline Bioluminescence Imaging counts and body weight. Mice were administered vehicle or Pegenzileukin (Sanofi) intravenously at 6 mg/kg at days 9, 16 and 23. At day 10, 2 × 106 CAR-positive T-cells or 2×106 untransduced T-cells were injected intravenously via the tail vein injection to selected groups. Mice were bled at specific intervals (days 14, 21, and 28), as per Sanofi IACUC guideline, to measure T-cell and CAR T-cell frequency and/or phenotype.

#### In-vivo Bioluminescence Imaging

Mice with Raji-Luc engraftment were anesthetized with isoflurane and were administered with D-luciferin (150 mg/kg) dissolved in PBS without calcium chloride and magnesium chloride (Perkin Elmer) through intraperitoneal injection. After the substrate injection, mice were placed into the IVIS imaging chamber stage (maintained at 37C) and imaged every 2 minutes up to 16 minutes. The imaging kinetics curves were obtained for each mouse, and peak imaging value was selected. A pseudocolor scheme was used to visualize the numerical contents of the acquired bioluminescent signal (superimposed onto contrast, gray-scale photographic pictures to determine the bioluminescence signal location). For whole body bioluminescence quantification, regions of interest (ROIs) were quantified as average radiance (photons/[s cm2 sr]), and the imaging signal graphic output was listed as total flux (photons s−1) of the whole body ROI.

#### Flow analysis

To detect T-cells in blood, 50ul blood sample ware lysed with 2ml RBC lysis buffer (Miltenyi Biotec) followed by centrifugation at 500×g for 10min. Cells were stained with live/dead staining using Zombie NIR Fixable Viability kit (Biolegend) followed by both human and mouse Fc blocking using Human TruStain FcX™ and anti-mouse CD16/32 TruStain FcX™ antibody, respectively (Biolegend). The cells were further stained with an antibody mix comprised of anti-mouse CD45 antibody and anti-human antibodies to the following: CD45, CD3, CD4, CD8, CD20 and as well as to human CD19 antigen. The stained samples were washed with FACS buffer comprised of 0.5% BSA (Sigma), PBS (Gibco) and 0.1% sodium azide (VWR Chemicals). The samples were recorded using a BD FACS Canto and data analysis was performed on FlowJo v10.9 software.

#### Histology and immunohistochemical staining

At study end, lung (inflated), liver, and spleen were fixed in 10% neutral buffered formalin for 48-72 hours, routinely processed into paraffin blocks, and sectioned at 5 microns. Standard staining using hematoxylin and eosin (H&E) was performed. Serial unstained sections were subjected to immunohistochemistry (IHC) using a rabbit anti-CD20 antibody (Therm Fisher Scientific) on BondRX Fully Automated Research Stainer (Leica Biosystems). H&E-stained slides and CD20 IHC slides were blindly evaluated by a board-certified veterinary pathologist. CD20 positive cell quantitation was performed on whole slide digital images using Cytonuclear (Lung) and Random Forest (Liver) algorithms of HALO image analysis software (Indica Labs).

## Supplemental Figure Legends

**Supplemental Figure 1:**
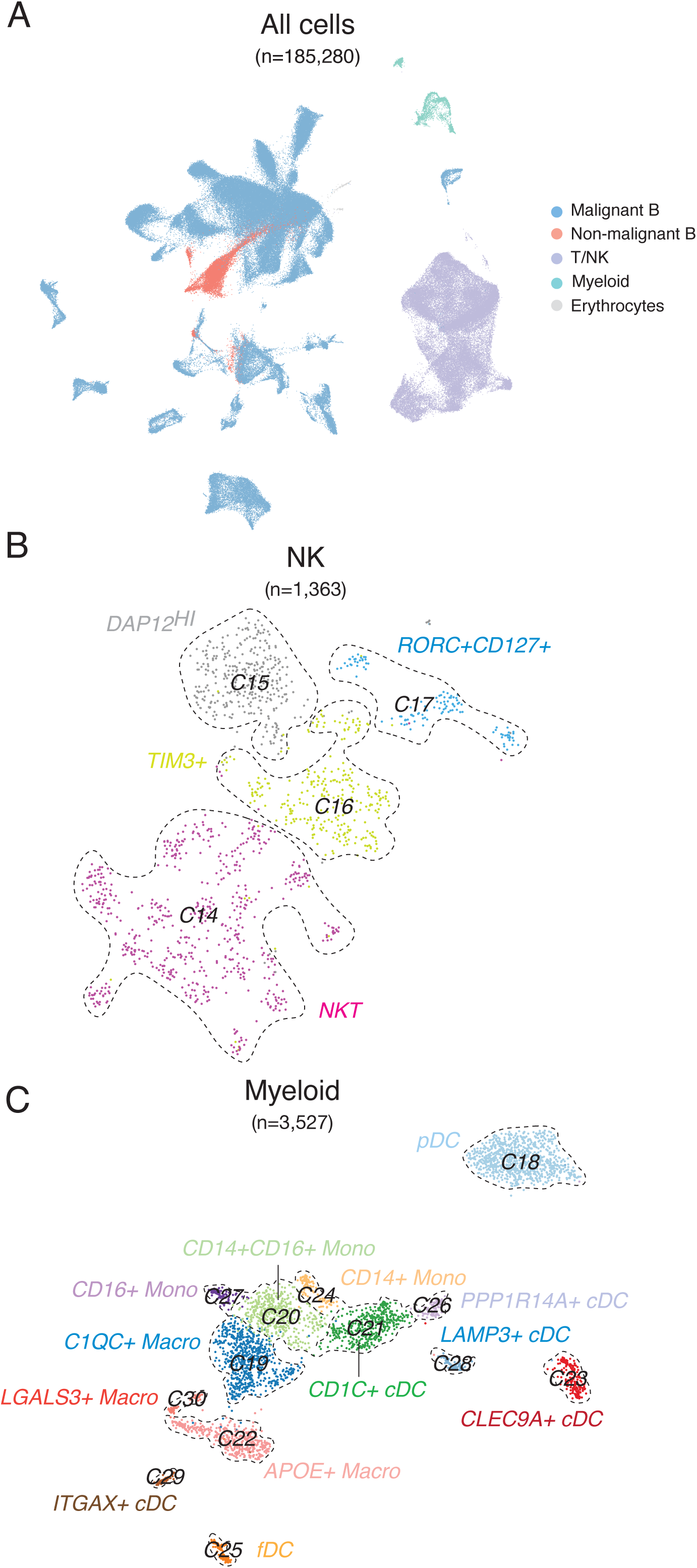
scRNA-seq Overall, NK, and Myeloid Cell Clusters. **A**. All sequenced single cells passing quality control (n = 185,280 cells) and cluster identities for malignant B cells, non-malignant B cells, T/NK cells, Myeloid cells, and Erythrocytes. **B.** Four clusters of NK cells (n = 1,365 cells). **C.** Thirteen clusters of myeloid cells (n = 3,527 cells).

**Supplemental Figure 2:**
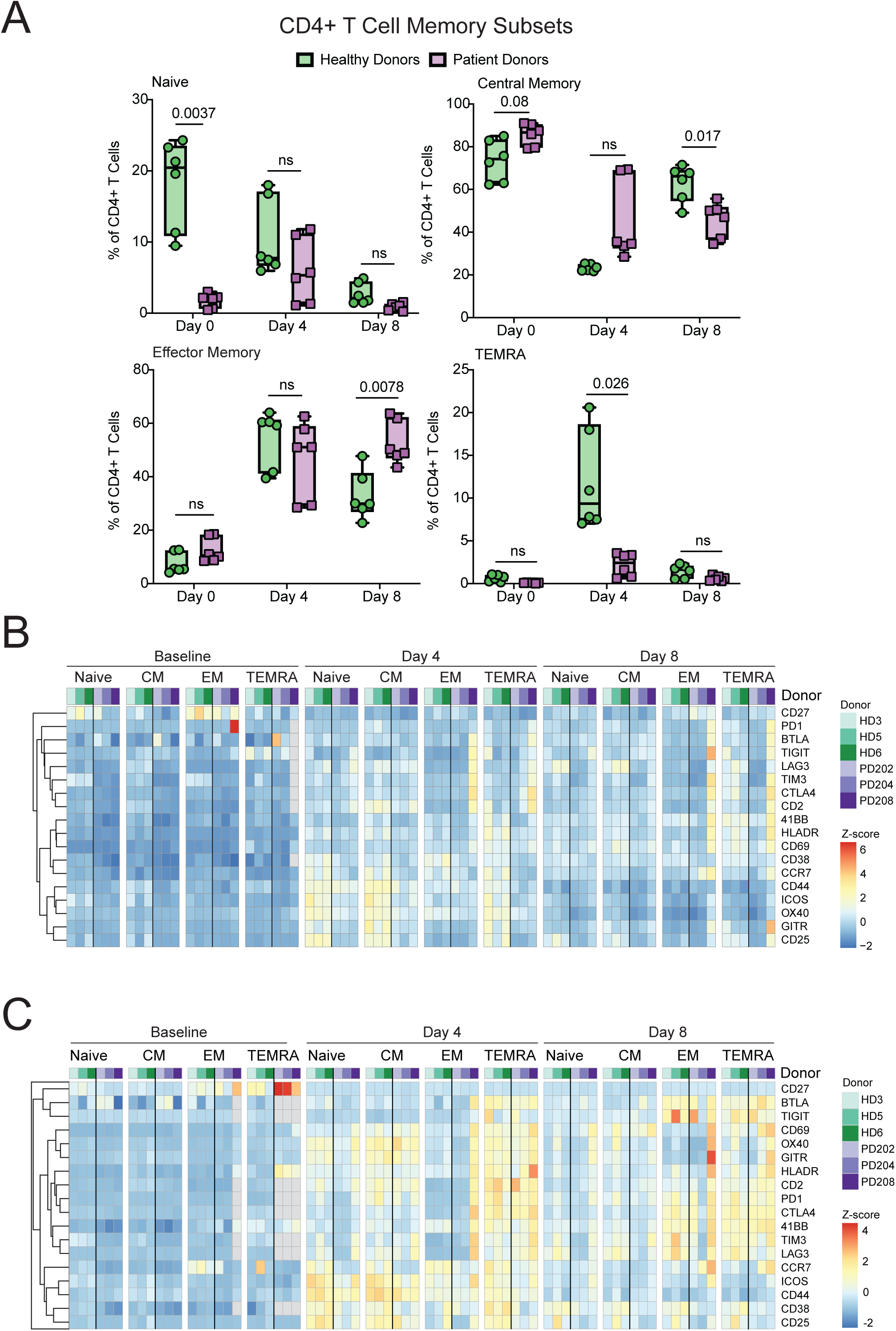
Healthy Donor Compared with Patient Derived T Cell Phenotypes. **A**. Changes in CD4 memory T cell phenotypes during serial re-challenge assay. Naïve (CD45RA+CD62L+), effector memory expressing CD45RA (TEMRA) (CD45RA+CD62L-), central memory (CD45RA-CD62L+), and effector memory (CD45RA-CD62L-). P-values shown are Bonferroni adjusted where adjusted p>0.05 = ns. **B.** Heatmap of CD8+ T cell surface immunophenotypes including grouping by day in tumor re-challenge assay, CD8 subsets, individual donors, and donor type. **C.** Heatmap of scaled MFI values of co-stimulatory and co-inhibitory markers within CD4 T cell memory subsets at each timepoint in serial re-challenge assay.

**Supplemental Figure 3:**
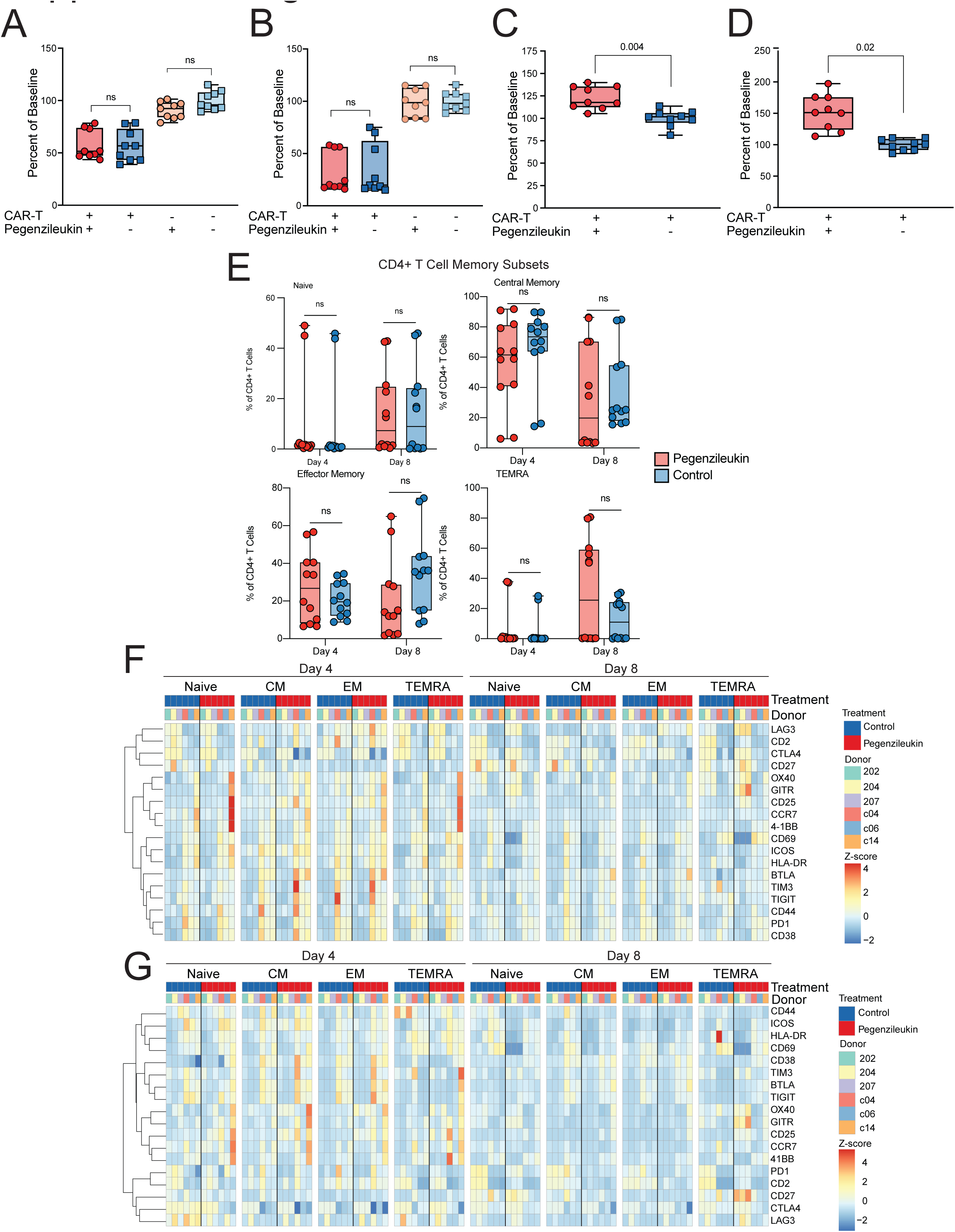
Pegenzileukin effects on CAR-T function and phenotype. **A**. 24-hour cytotoxicity at 1:1 effector to target ratio of RL LBCL cells with or without CD19 CAR T cells and with or without addition of pegenzileukin. Data presented as lymphoma cell count as a percentage of baseline. **B.** T cell proliferation over 24 hours with or without addition of pegenzileukin. **C.** 48-hour cytotoxicity at 1:1 effector to target ratio of RL LBCL cells with or without CD19 CAR T cells and with or without addition of pegenzileukin. Data presented as lymphoma cell count as a percentage of baseline. **D.** T cell proliferation over 48 hours with or without addition of pegenzileukin. **E.** Changes in CD4 memory T cell phenotypes during serial re-challenge assay. Naïve (CD45RA+CD62L+), effector memory expressing CD45RA (TEMRA) (CD45RA+CD62L-), central memory (CD45RA-CD62L+), and effector memory (CD45RA-CD62L-). P-values shown are Bonferroni adjusted where adjusted p>0.05 = ns. **F.** Heatmap of CD8+ T cell surface immunophenotypes including grouping by day in tumor re-challenge assay, CD8 subsets, treatment with control or pegenzileukin and individual patient donors. **G.** Heatmap of scaled MFI values of co-stimulatory and co-inhibitory markers within CD4 T cell memory subsets at each timepoint in serial re-challenge assay.

## REFERENCES

1. Locke, F.L., Miklos, D.B., Jacobson, C.A., Perales, M.-A., Kersten, M.-J., Oluwole, O.O., Ghobadi, A., Rapoport, A.P., McGuirk, J., Pagel, J.M., et al. (2021). Axicabtagene Ciloleucel as Second-Line Therapy for Large B-Cell Lymphoma. New England Journal of Medicine 386, 640–654. 10.1056/NEJMoa2116133.

2. Spiegel, J.Y., Dahiya, S., Jain, M.D., Tamaresis, J., Nastoupil, L.J., Jacobs, M.T., Ghobadi, A., Lin, Y., Lunning, M., Lekakis, L., et al. (2021). Outcomes of patients with large B-cell lymphoma progressing after axicabtagene ciloleucel therapy. Blood 137, 1832–1835. 10.1182/blood.2020006245.

3. Ayuk, F.A., Berger, C., Badbaran, A., Zabelina, T., Sonntag, T., Riecken, K., Geffken, M., Wichmann, D., Frenzel, C., Thayssen, G., et al. (2021). Axicabtagene ciloleucel in vivo expansion and treatment outcome in aggressive B-cell lymphoma in a real-world setting. Blood Adv 5, 2523–2527. 10.1182/bloodadvances.2020003959.

4. Locke, F.L., Rossi, J.M., Neelapu, S.S., Jacobson, C.A., Miklos, D.B., Ghobadi, A., Oluwole, O.O., Reagan, P.M., Lekakis, L.J., Lin, Y., et al. (2020). Tumor burden, inflammation, and product attributes determine outcomes of axicabtagene ciloleucel in large B-cell lymphoma. Blood Adv 4, 4898–4911. 10.1182/bloodadvances.2020002394.

5. Monfrini, C., Stella, F., Aragona, V., Magni, M., Ljevar, S., Vella, C., Fardella, E., Chiappella, A., Nanetti, F., Pennisi, M., et al. (2022). Phenotypic Composition of Commercial Anti-CD19 CAR T Cells Affects In Vivo Expansion and Disease Response in Patients with Large B-cell Lymphoma. Clinical cancer research: an official journal of the American Association for Cancer Research 28, 3378–3386. 10.1158/1078-0432.CCR-22-0164.

6. Deng, Q., Han, G., Puebla-Osorio, N., Ma, M.C.J., Strati, P., Chasen, B., Dai, E., Dang, M., Jain, N., Yang, H., et al. (2020). Characteristics of anti-CD19 CAR T cell infusion products associated with efficacy and toxicity in patients with large B cell lymphomas. Nat Med 26, 1878–1887. 10.1038/s41591-020-1061-7.

7. Fuller, M.J., and Zajac, A.J. (2003). Ablation of CD8 and CD4 T cell responses by high viral loads. J Immunol 170, 477–486. 10.4049/jimmunol.170.1.477.

8. Wherry, E.J. (2011). T cell exhaustion. Nature immunology 12, 492–499. 10.1038/ni.2035.

9. Wherry, E.J., Blattman, J.N., Murali-Krishna, K., van der Most, R., and Ahmed, R. (2003). Viral persistence alters CD8 T-cell immunodominance and tissue distribution and results in distinct stages of functional impairment. J Virol 77, 4911–4927. 10.1128/jvi.77.8.4911-4927.2003.

10. Chauvin, J.M., Pagliano, O., Fourcade, J., Sun, Z., Wang, H., Sander, C., Kirkwood, J.M., Chen, T.H., Maurer, M., Korman, A.J., and Zarour, H.M. (2015). TIGIT and PD-1 impair tumor antigen-specific CD8⁺ T cells in melanoma patients. J Clin Invest 125, 2046–2058. 10.1172/jci80445.

11. Zhang, Z., Liu, S., Zhang, B., Qiao, L., Zhang, Y., and Zhang, Y. (2020). T Cell Dysfunction and Exhaustion in Cancer. Frontiers in Cell and Developmental Biology 8. 10.3389/fcell.2020.00017.

12. Rosenberg, S.A., Packard, B.S., Aebersold, P.M., Solomon, D., Topalian, S.L., Toy, S.T., Simon, P., Lotze, M.T., Yang, J.C., Seipp, C.A., et al. (1988). Use of Tumor-Infiltrating Lymphocytes and Interleukin-2 in the Immunotherapy of Patients with Metastatic Melanoma. New England Journal of Medicine 319, 1676–1680. 10.1056/nejm198812223192527.

13. Kalia, V., and Sarkar, S. (2018). Regulation of Effector and Memory CD8 T Cell Differentiation by IL-2-A Balancing Act. Front Immunol 9, 2987. 10.3389/fimmu.2018.02987.

14. Zheng, S.G., Wang, J., Wang, P., Gray, J.D., and Horwitz, D.A. (2007). IL-2 Is Essential for TGF-β to Convert Naive CD4+CD25− Cells to CD25+Foxp3+ Regulatory T Cells and for Expansion of These Cells1. The Journal of Immunology 178, 2018–2027. 10.4049/jimmunol.178.4.2018.

15. Krieg, C., Létourneau, S., Pantaleo, G., and Boyman, O. (2010). Improved IL-2 immunotherapy by selective stimulation of IL-2 receptors on lymphocytes and endothelial cells. Proceedings of the National Academy of Sciences 107, 11906–11911. doi:10.1073/pnas.1002569107.

16. Van Gool, F., Molofsky, A.B., Morar, M.M., Rosenzwajg, M., Liang, H.E., Klatzmann, D., Locksley, R.M., and Bluestone, J.A. (2014). Interleukin-5-producing group 2 innate lymphoid cells control eosinophilia induced by interleukin-2 therapy. Blood 124, 3572–3576. 10.1182/blood-2014-07-587493.

17. Good, Z., Spiegel, J.Y., Sahaf, B., Malipatlolla, M.B., Ehlinger, Z.J., Kurra, S., Desai, M.H., Reynolds, W.D., Wong Lin, A., Vandris, P., et al. (2022). Post-infusion CAR TReg cells identify patients resistant to CD19-CAR therapy. Nature Medicine 28, 1860–1871. 10.1038/s41591-022-01960-7.

18. Haradhvala, N.J., Leick, M.B., Maurer, K., Gohil, S.H., Larson, R.C., Yao, N., Gallagher, K.M.E., Katsis, K., Frigault, M.J., Southard, J., et al. (2022). Distinct cellular dynamics associated with response to CAR-T therapy for refractory B cell lymphoma. Nature Medicine 28, 1848–1859. 10.1038/s41591-022-01959-0.

19. Ptacin, J.L., Caffaro, C.E., Ma, L., San Jose Gall, K.M., Aerni, H.R., Acuff, N.V., Herman, R.W., Pavlova, Y., Pena, M.J., Chen, D.B., et al. (2021). An engineered IL-2 reprogrammed for anti-tumor therapy using a semi-synthetic organism. Nature Communications 12, 4785. 10.1038/s41467-021-24987-9.

20. Han, G., Deng, Q., Marques-Piubelli, M.L., Dai, E., Dang, M., Ma, M.C.J., Li, X., Yang, H., Henderson, J., Kudryashova, O., et al. (2022). Follicular Lymphoma Microenvironment Characteristics Associated with Tumor Cell Mutations and MHC Class II Expression. Blood Cancer Discov 3, 428–443. 10.1158/2643-3230.Bcd-21-0075.

21. DeRogatis, J.M., Neubert, E.N., Viramontes, K.M., Henriquez, M.L., Nicholas, D.A., and Tinoco, R. (2023). Cell-Intrinsic CD38 Expression Sustains Exhausted CD8(+) T Cells by Regulating Their Survival and Metabolism during Chronic Viral Infection. J Virol 97, e0022523. 10.1128/jvi.00225-23.

22. Rutishauser, R.L., Martins, G.A., Kalachikov, S., Chandele, A., Parish, I.A., Meffre, E., Jacob, J., Calame, K., and Kaech, S.M. (2009). Transcriptional repressor Blimp-1 promotes CD8(+) T cell terminal differentiation and represses the acquisition of central memory T cell properties. Immunity 31, 296–308. 10.1016/j.immuni.2009.05.014.

23. Zhang, X., Zhang, C., Qiao, M., Cheng, C., Tang, N., Lu, S., Sun, W., Xu, B., Cao, Y., Wei, X., et al. (2022). Depletion of BATF in CAR-T cells enhances antitumor activity by inducing resistance against exhaustion and formation of central memory cells. Cancer Cell 40, 1407–1422.e1407. 10.1016/j.ccell.2022.09.013.

24. Chu, Y., Dai, E., Li, Y., Han, G., Pei, G., Ingram, D.R., Thakkar, K., Qin, J.J., Dang, M., Le, X., et al. (2023). Pan-cancer T cell atlas links a cellular stress response state to immunotherapy resistance. Nat Med 29, 1550–1562. 10.1038/s41591-023-02371-y.

25. Carrio, R., Cucchetti, M., Devonish, M., Poisson, L., Babiceanu, M., Kettring, A., Liu, Y., Jackson, D., Gomes, E., Baudhuin, J., et al. (2023). Abstract 6378: An in vitro human CD8 T cell exhaustion model for the functional screening of immune checkpoint inhibitors. Cancer research 83, 6378–6378. 10.1158/1538-7445.Am2023-6378.

26. Singh, N., Lee, Y.G., Shestova, O., Ravikumar, P., Hayer, K.E., Hong, S.J., Lu, X.M., Pajarillo, R., Agarwal, S., Kuramitsu, S., et al. (2020). Impaired Death Receptor Signaling in Leukemia Causes Antigen-Independent Resistance by Inducing CAR T-cell Dysfunction. Cancer discovery 10, 552–567. 10.1158/2159-8290.CD-19-0813.

27. Decker, C.E., Young, T., Pasnikowski, E., Chiu, J., Song, H., Wei, Y., Thurston, G., and Daly, C. (2019). Genome-scale CRISPR activation screen uncovers tumor-intrinsic modulators of CD3 bispecific antibody efficacy. Sci Rep 9, 20068. 10.1038/s41598-019-56670-x.

28. Liu, Y., Zhou, N., Zhou, L., Wang, J., Zhou, Y., Zhang, T., Fang, Y., Deng, J., Gao, Y., Liang, X., et al. (2021). IL-2 regulates tumor-reactive CD8(+) T cell exhaustion by activating the aryl hydrocarbon receptor. Nat Immunol 22, 358–369. 10.1038/s41590-020-00850-9.

29. Finney, O.C., Brakke, H.M., Rawlings-Rhea, S., Hicks, R., Doolittle, D., Lopez, M., Futrell, R.B., Orentas, R.J., Li, D., Gardner, R.A., and Jensen, M.C. (2019). CD19 CAR T cell product and disease attributes predict leukemia remission durability. J Clin Invest 129, 2123–2132. 10.1172/jci125423.

30. Fraietta, J.A., Lacey, S.F., Orlando, E.J., Pruteanu-Malinici, I., Gohil, M., Lundh, S., Boesteanu, A.C., Wang, Y., O’Connor, R.S., Hwang, W.T., et al. (2018). Determinants of response and resistance to CD19 chimeric antigen receptor (CAR) T cell therapy of chronic lymphocytic leukemia. Nature medicine 24, 563–571. 10.1038/s41591-018-0010-1.

31. Jain, M.D., Zhao, H., Wang, X., Atkins, R., Menges, M., Reid, K., Spitler, K., Faramand, R., Bachmeier, C., Dean, E.A., et al. (2021). Tumor interferon signaling and suppressive myeloid cells are associated with CAR T-cell failure in large B-cell lymphoma. Blood 137, 2621–2633. 10.1182/blood.2020007445.

32. Scholler, N., Perbost, R., Locke, F.L., Jain, M.D., Turcan, S., Danan, C., Chang, E.C., Neelapu, S.S., Miklos, D.B., Jacobson, C.A., et al. (2022). Tumor immune contexture is a determinant of anti-CD19 CAR T cell efficacy in large B cell lymphoma. Nature Medicine 28, 1872–1882. 10.1038/s41591-022-01916-x.

33. Gumber, D., and Wang, L.D. (2022). Improving CAR-T immunotherapy: Overcoming the challenges of T cell exhaustion. EBioMedicine 77, 103941. 10.1016/j.ebiom.2022.103941.

34. Poorebrahim, M., Melief, J., Pico de Coaña, Y.S.L.W., Cid-Arregui, A., and Kiessling, R. (2021). Counteracting CAR T cell dysfunction. Oncogene 40, 421–435. 10.1038/s41388-020-01501-x.

35. Hinrichs, C.S., Borman, Z.A., Gattinoni, L., Yu, Z., Burns, W.R., Huang, J., Klebanoff, C.A., Johnson, L.A., Kerkar, S.P., Yang, S., et al. (2011). Human effector CD8+ T cells derived from naive rather than memory subsets possess superior traits for adoptive immunotherapy. Blood 117, 808–814. 10.1182/blood-2010-05-286286.

36. Zhang, D.K.Y., Adu-Berchie, K., Iyer, S., Liu, Y., Cieri, N., Brockman, J.M., Neuberg, D., Wu, C.J., and Mooney, D.J. (2023). Enhancing CAR-T cell functionality in a patient-specific manner. Nature Communications 14, 506. 10.1038/s41467-023-36126-7.

37. Janku, F., Abdul-Karim, R., Azad, A., Bendell, J., Falchook, G., Gan, H.K., Tan, T., Wang, J.S., Chee, C.E., Ma, L., et al. (2021). Abstract LB041: THOR-707 (SAR444245), a novel not-alpha IL-2 as monotherapy and in combination with pembrolizumab in advanced/metastatic solid tumors: Interim results from HAMMER, an open-label, multicenter phase 1/2 Study. Cancer Research 81, LB041–LB041. 10.1158/1538-7445.Am2021-lb041.

38. Cherng, H.J., Sun, R., Sugg, B., Irwin, R., Yang, H., Le, C.C., Deng, Q., Fayad, L., Fowler, N.H., Parmar, S., et al. (2022). Risk assessment with low-pass whole-genome sequencing of cell-free DNA before CD19 CAR T-cell therapy for large B-cell lymphoma. Blood 140, 504–515. 10.1182/blood.2022015601.

39. Jain, M.D., Ziccheddu, B., Coughlin, C.A., Faramand, R., Griswold, A.J., Reid, K.M., Menges, M., Zhang, Y., Cen, L., Wang, X., et al. (2022). Whole-genome sequencing reveals complex genomic features underlying anti-CD19 CAR T-cell treatment failures in lymphoma. Blood 140, 491–503. 10.1182/blood.2021015008.

40. Jacobson, C.A., Hunter, B.D., Redd, R., Rodig, S.J., Chen, P.H., Wright, K., Lipschitz, M., Ritz, J., Kamihara, Y., Armand, P., et al. (2020). Axicabtagene Ciloleucel in the Non-Trial Setting: Outcomes and Correlates of Response, Resistance, and Toxicity. J Clin Oncol 38, 3095–3106. 10.1200/jco.19.02103.

41. Nastoupil, L.J., Jain, M.D., Feng, L., Spiegel, J.Y., Ghobadi, A., Lin, Y., Dahiya, S., Lunning, M., Lekakis, L., Reagan, P., et al. (2020). Standard-of-Care Axicabtagene Ciloleucel for Relapsed or Refractory Large B-Cell Lymphoma: Results From the US Lymphoma CAR T Consortium. J Clin Oncol 38, 3119–3128. 10.1200/JCO.19.02104.

42. Bethge, W.A., Martus, P., Schmitt, M., Holtick, U., Subklewe, M., von Tresckow, B., Ayuk, F., Wagner-Drouet, E.M., Wulf, G.G., Marks, R., et al. (2022). GLA/DRST real-world outcome analysis of CAR T-cell therapies for large B-cell lymphoma in Germany. Blood 140, 349–358. 10.1182/blood.2021015209.

43. Dean, E.A., Mhaskar, R.S., Lu, H., Mousa, M.S., Krivenko, G.S., Lazaryan, A., Bachmeier, C.A., Chavez, J.C., Nishihori, T., Davila, M.L., et al. (2020). High metabolic tumor volume is associated with decreased efficacy of axicabtagene ciloleucel in large B-cell lymphoma. Blood Adv 4, 3268–3276. 10.1182/bloodadvances.2020001900.

